# New tools for diet analysis: nanopore sequencing of metagenomic DNA from rat stomach contents to quantify diet

**DOI:** 10.1101/363622

**Authors:** Nikki E. Freed, William S. Pearman, Adam N. H. Smith, Georgia Breckell, James Dale, Olin K. Silander

**Affiliations:** Institute of Natural and Mathematical Sciences, Massey University, Auckland 0745, New Zealand

**Author notes:** Corresponding authors: Olin K. Silander, Institute of Natural and Mathematical Sciences, Massey University, Auckland 0745, New Zealand; Nikki E. Freed, Institute of Natural and Mathematical Sciences, Massey University, Auckland 0745, New Zealand.

**Keywords:** Metagenomics, nanopore, diet analysis, rat, next generation sequencing

## Abstract

**Background:** Using metagenomics to determine animal diet offers a new and promising alternative to current methods. Here we show that rapid and inexpensive diet quantification is possible through metagenomic sequencing with the portable Oxford Nanopore Technologies (ONT) MinION. Using an amplification-free approach, we profiled the stomach contents from wild-caught rats.

**Results:** We conservatively identified diet items from over 50 taxonomic orders, ranging across nine phyla that include plants, vertebrates, invertebrates, and fungi. This highlights the wide range of taxa that can be identified using this simple approach. We calibrate the accuracy of this method by comparing the characteristics of reads matching the ground-truth host genome (rat) to those matching diet items, and show that at the family-level, false positive taxon assignments are approximately 97.5% accurate. We also suggest a way to mitigate for database biases in metagenomic approaches. Finally, we implement a constrained ordination analysis and show that we can identify the sampling location of an individual rat within tens of kilometres based on diet content alone.

**Conclusions:** This work establishes proof-of-principle for long-read metagenomic methods in quantitative diet analysis. We show that diet content can be quantified even with limited expertise, using a simple, amplification free workflow and a relatively inexpensive and accessible next generation sequencing method. Continued increases in the accuracy and throughput of ONT sequencing, along with improved genomic databases, suggests that a metagenomic approach to quantification of animal diets will become an important method in the future.

## Background

Accurate quantification of animal diets yields critical insight into ecosystem and food web dynamics. However, unbiased and sensitive assessment of diet content is difficult to achieve. This is largely due to the limited accuracy of many current methods. Such methods include visual inspection of gut contents (1,2), which presents bias against items are most easily degraded (for example, soft-bodied species); stable isotope analysis (3,4), which yields only broad information on diet, such as whether diet items are terrestrial or marine in origin (5,6); and time-lapse video (7,8), for which species identification is difficult for small prey items or in low-light conditions.

To circumvent these issues, DNA-based methods (9,10) have become popular. Perhaps the most widely applied DNA-based method is metabarcoding. This approach relies on PCR amplification and sequencing of conserved regions from nuclear, mitochondrial, or plastid genomes (9). With adequate primer selection, this method can detect a wide range of species, and does not require specific expertise, which is often necessary for other methods.

However, DNA metabarcoding is not free from bias. PCR primers must be specifically tailored to particular sets of taxa or species (11). Although “universal” PCR primer pairs have been developed (for example targeting all bilaterians or even all eukaryotes (12), all primer sets exhibit bias towards certain taxa. Five-fold differences in fungal operational taxonomic units (OTU) estimates have been found when using different sets of fungal-specific PCR primer pairs (13). It has also been shown that published universal primer pairs were capable of amplifying only between 57% and 91% of tested metazoan species, with as few as 33% of species in some phyla being amplified at all (e.g. cnidarians)(14). Different genomic loci from the same species can exhibit up to 2,000-fold differences in DNA concentration as inferred using qPCR (15). The choice of polymerase can also bias diversity metrics when using metabarcoding (16). For these reasons, an approach that circumvents PCR and thus avoids these biases is desirable.

Metagenomic sequencing aims to directly sequence all of the DNA in a sample without introducing bias. Although there are still biases with this approach, for example due to nucleotide content affecting the likelihood of a molecule being sequenced, these are inherently less than those introduced by metabarcoding. Metagenomic approaches have most frequently been used to yield insights into microbial diversity and function (17–24). Metagenomic applications aimed at eukaryotic taxa identification are less common. Several metagenomic diet studies have implemented filtering steps to select only mitogenomic sequence or metabarcode regions, or have used abridged databases before data analysis in order to mitigate database biases (25–27). However, to our knowledge, very few studies have used unfiltered metagenomic sequence analysis to infer diet (30,31).

Here, we establish a proof-of-principle methodology to accurately classify metagenomic sequences from eukaryotic taxa and determine diet content using low-accuracy, long-read sequencing, Oxford Nanopore (ONT).

We quantified rat diets from several locations in the North Island of New Zealand using stomach samples. Using these samples and methodology provides three distinct advantages.

First, rats are extremely omnivorous. As such, they serve as an excellent means to quantify the breadth of taxa that can be detected using a metagenomic long read approach.

Second, the use of stomach samples means that a significant number of reads will be host reads. This allows us to assess the characteristics of true positive sequence reads (rat-derived reads that match rat database sequences), as well as false positive reads (rat-derived reads that match non-rat database sequences). We can then determine whether reads matching diet items have similar characteristics to known true positive reads. This use of host reads is exactly analogous to feeding the rats a diet of known content (i.e. rat) and testing whether the contents of the known diet can be accurately identified using this method.

Third, quantifying rat diets has important ecological implications. It is well-established that the relatively recent introduction of mammalian predators to New Zealand has had significant negative effects on many of the native animal populations. This ranges from insects (34), to reptiles (35), to molluscs (36), to birds (37,38), with downstream effects on terrestrial and aquatic ecosystems (39). To counteract the effects of mammalian predators, an ambitious plan is currently being put into place that aims for the eradication of all mammalian predators from New Zealand (including possums, rats, stoats, and hedgehogs), by 2050 (http://www.doc.govt.nz/predator-free-2050; (40). A useful step toward this goal would be to prioritise the management of predators and establish in which locations native species experience the highest levels of predation. To do so requires establishing the diet content of local mammalian predators.

Importantly, using DNA sequencing methods to profile diet does not require high depth to accurately profile the breadth of diet items consumed by an animal. Here we aim to quantify all diet items present in the diet at 1% or more. At this fraction, if we assume that read counts are Poisson distributed, with only 2,000 reads, 99% of the time we will quantify such items within 2-fold of their true amount. Thus, diet quantification presents a clear example of a situation in which sequencing depth is not a critical factor.

## Results

### DNA sequencing and assignment of reads to taxa

We selected eight rats from each of three locations near Auckland, New Zealand for diet quantification. Each location comprised a different type of habitat: undisturbed inland native forest (Waitakere Regional Parklands, WP); native bush surrounding an estuary (Okura Bush Walkway, OB); and restored coastal wetland (Long Bay Regional Park, LB). We isolated DNA from whole stomach contents from each rat (see Methods). We obtained DNA from entire homogenised stomach samples. We sequenced these DNA samples using the Oxford Nanopore ligation sequencing kit (LSK-108) and native barcoding kit to multiplex the samples within a sequencing run. We performed two separate runs, obtaining a total of 82,977 reads (January 2017) and 96,150 reads (March 2017) from each. These numbers are not far below expectations given the flow cell and kit chemistry and MinKNOW software versions available at that time (47). However, these read numbers are considerably below those expected for current ONT flow cells and software, which has improved per flow cell output more than 100-fold above these numbers.

We found large variation in the numbers of reads per barcode: approximately 10-fold for the January samples, and up to 40-fold in March (Fig. 1A and 1B). We hypothesise that this is due to the highly variable quality of DNA in each sample. This did not appear to have strong effects on read accuracy, as the median phred quality scores per read ranged from 7-12 (0.80 - 0.94 accuracy) for both runs.

**Fig. 1.**
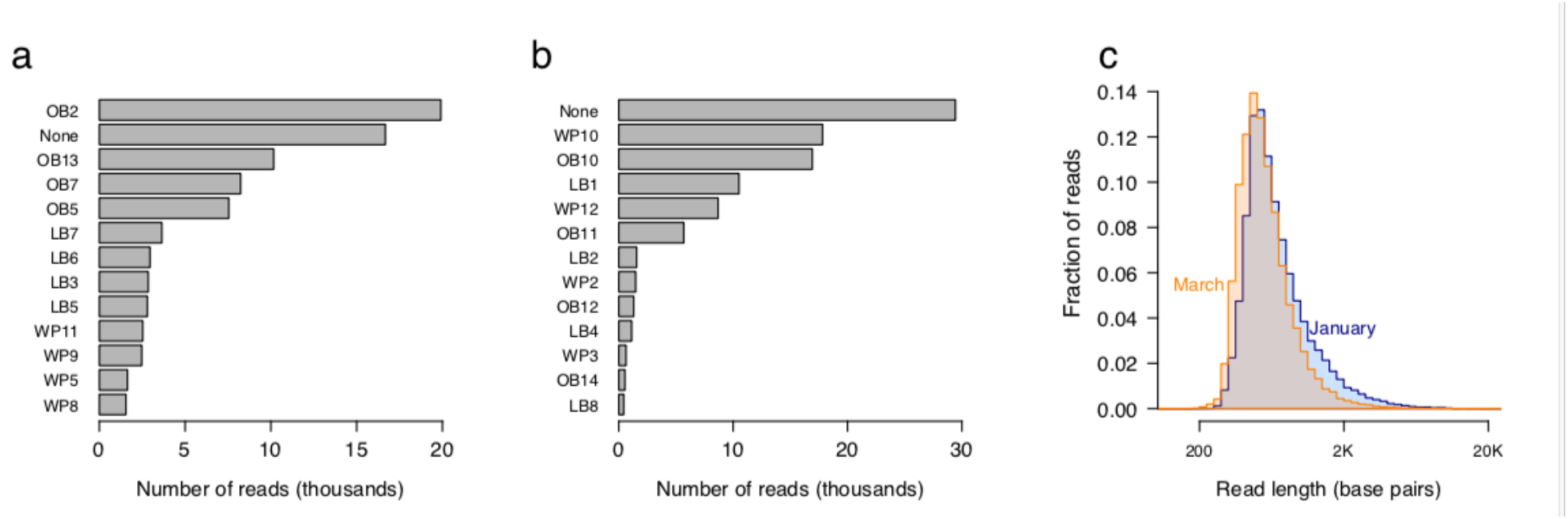
Results of nanopore metagenomic sequencing of rat stomach contents. (A) and (B) Barcode distributions for January and March runs, respectively. We multiplexed the samples on the flow cells, using 12 barcodes per flow cell. The distribution of read numbers across barcodes varied by up to 40-fold. 20% (January) and 30% (March) of all reads could not be assigned to a barcode (“None”). The inability to assign these reads to a barcode is due primarily to their lower quality. **(C) Read length distribution for January and March nanopore runs.** 90% of the reads were between 350 bp and 1,580 bp in length, with only 0.55% being longer than 4,000 bp.

The degradation of the DNA during digestion in the stomach, as well as fragmentation during DNA isolation (48) and sequencing library preparation led to relatively short median read lengths of 606 bp and 527 bp for the January and March datasets, respectively (Fig. 1C). Within runs, read length and quality were similar. Although these median read lengths are considerably shorter than other nanopore sequencing results from both our and others work (49), we found wide variation in length, with almost 10% of all reads being longer than 1200 bp. Notably, reads of this length can allow more precise taxonomic identification than accurate short reads (32).

To quantify diet contents we first BLASTed all sequences against a combined database of the NCBI nt database (the partially non-redundant nucleotide sequences from all traditional divisions of GenBank excluding genome survey sequence, EST, high-throughput genome, and whole genome shotgun (ftp://ftp.ncbi.nlm.nih.gov/blast/db/README)) and the NCBI other_genomic database (RefSeq chromosome records for non-human organisms (ftp://ftp.ncbi.nlm.nih.gov/blast/db/README)). We used BLAST as it is generally viewed as the gold standard method in metagenomic analyses (50). Of the 133,022 barcoded reads, 30,535 (23%) hit a sequence in the combined nt and other_genomic database at an e-value cut-off of 1e-2.

We first aimed to assess the quality of these hits. We found a bimodal distribution of alignment lengths and e-values (Fig. 2A). We also noticed that mean read quality had substantial effects on the likelihood of a read yielding a BLAST hit, with almost 40% of high accuracy reads having hits in the March dataset, as compared to 1% of low accuracy reads (Fig. 2B). We hypothesized that many of the short alignments with high e-values were false positives. We thus first filtered this hit set, only retaining BLAST hits with e-values less than 1e-20 and alignments greater than 100 bp. To give some intuition for this length cut-off, a 100bp read with five 1 bp indels and seven mismatches (88% identity) would have an identical raw score and e-value (given the default match, mismatch, and gap parameters) to a single-end 70 bp Illumina read with a 100% identical database match. Similar or quality filters based on length and identity have been imposed previously (30). A total of 22,154 hits passed this e-value filter.

**Fig. 2.**
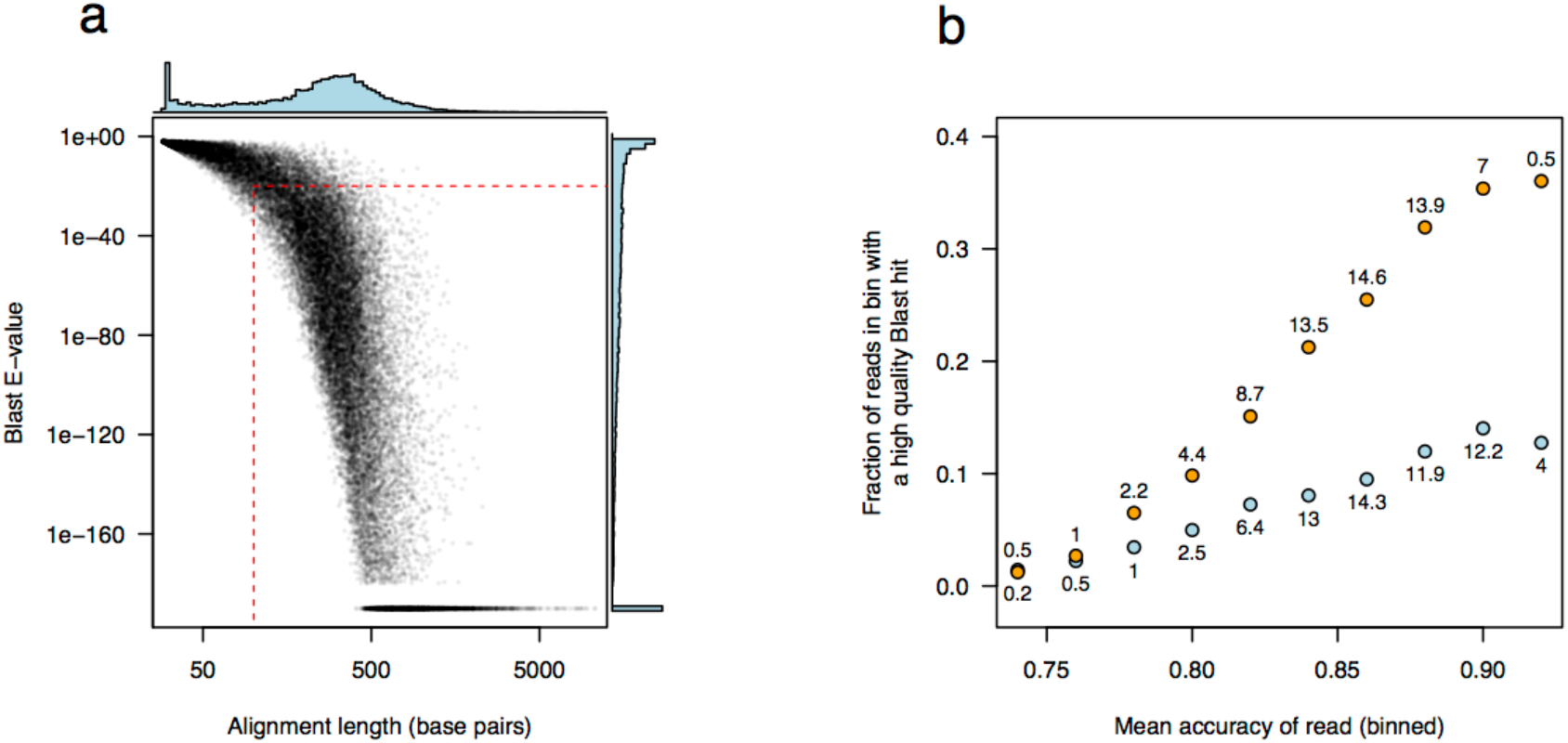
BLAST hits of metagenomic reads. **(a)** Alignment lengths and e-values were bimodally distributed. The y-axis is plotted on a log scale, with zero e-values suppressed by adding a small number (1e-190) to each e-value. The horizontal red dotted line indicates the e-value cut-off we implemented and the vertical red dotted line indicates the length cut-off (e-value < 1e-20 and alignment length of 100, respectively) to decrease false positive hits. **(b) The fraction of reads with high quality BLAST hits (e-value < 1e-20) increased as a function of read accuracy.** We binned the data according to mean read accuracy (bin width = 0.02) and calculated the fraction of reads within each bin that have a high quality BLAST hit (alignment length greater than 100bp and e-value less than 1e-20) for the January and March runs separately (blue and orange points, respectively). The number of reads in each bin is indicated above each point (in thousands). There is a clear positive correlation between mean accuracy and the likelihood of a high quality BLAST hit, reaching almost 40% for high accuracy reads (>92.5%) for the March dataset.

We next used MEGAN6 (41) to assign reads to specific taxa. MEGAN6 employs an LCA algorithm to assign reads to a taxon. For example, if a read has BLAST hits to five different species, three of which have bit scores within 20% of the best hit, the read will be assigned to the genus, family, order, or higher taxon level that is the LCA of those best-hit three species (51). If a read matches one species far better than any other, by definition, the LCA is that species.

16,820 reads (76%) were assigned to a taxon by MEGAN. Of these, 31% were assigned by MEGAN as being bacterial, and 55% of these were *Lactobacillus spp*. These results match previous studies on rat stomach microbiomes, which have found lactobacilli to be the dominant taxa (52–55). Plant-associated *Pseudomonas* and *Lactococcus* taxa were also common, at 7% and 6%, respectively.

MEGAN assigned reads to a wide range of eukaryotic taxa. To conservatively infer taxon presence, we first reclassified MEGAN species-level assignments to the level of genus. Even so, many clear false positive assignments remained (e.g. hippo and naked mole rat). These matches were generally short and of low identity. To reduce such false positive taxon inferences, we used information from reads assigned to the genera *Rattus* (rat) and *Mus* (mouse).

We inferred that the reads assigned to *Rattus* (2,696 reads in total) were true positive genus-level assignments and that the reads assigned to *Mus* (2,798 reads in total) were false positive genus-level assignments (and not true positive *Mus*-derived reads). If these were true positive *Mus* reads, then they would be due to mouse predation by some of the rats. Although rats are known to prey on mice (56), if this had occurred, we would expect that (1) if a rat predated a mouse the ratio of mouse to rat reads would be higher than in rats that had not predated mice; (2) in those same rats, the percent identity of the reads assigned to *Mus* would be higher than in rats that had not predated mice. However, we found that the ratio of mouse to rat reads and percent identity was similar for all rats. This suggested that these *Mus* hits were indeed false positives and could thus be used to determine the rate of false positive genus-level assignment.

The mean percent identity values of the best BLAST hits for *Rattus* and *Mus* reads differed substantially, with *Rattus* reads having a median identity of 86.4%, and *Mus* 81.0% (Fig. 3A). The mean percent identity for *Rattus* reads corresponds very well to that expected given the mean quality scores of the reads (86.4% identity corresponds to a mean quality score of 8.7, similar to what we observed; Fig. S1A-C).

**Fig. 3.**
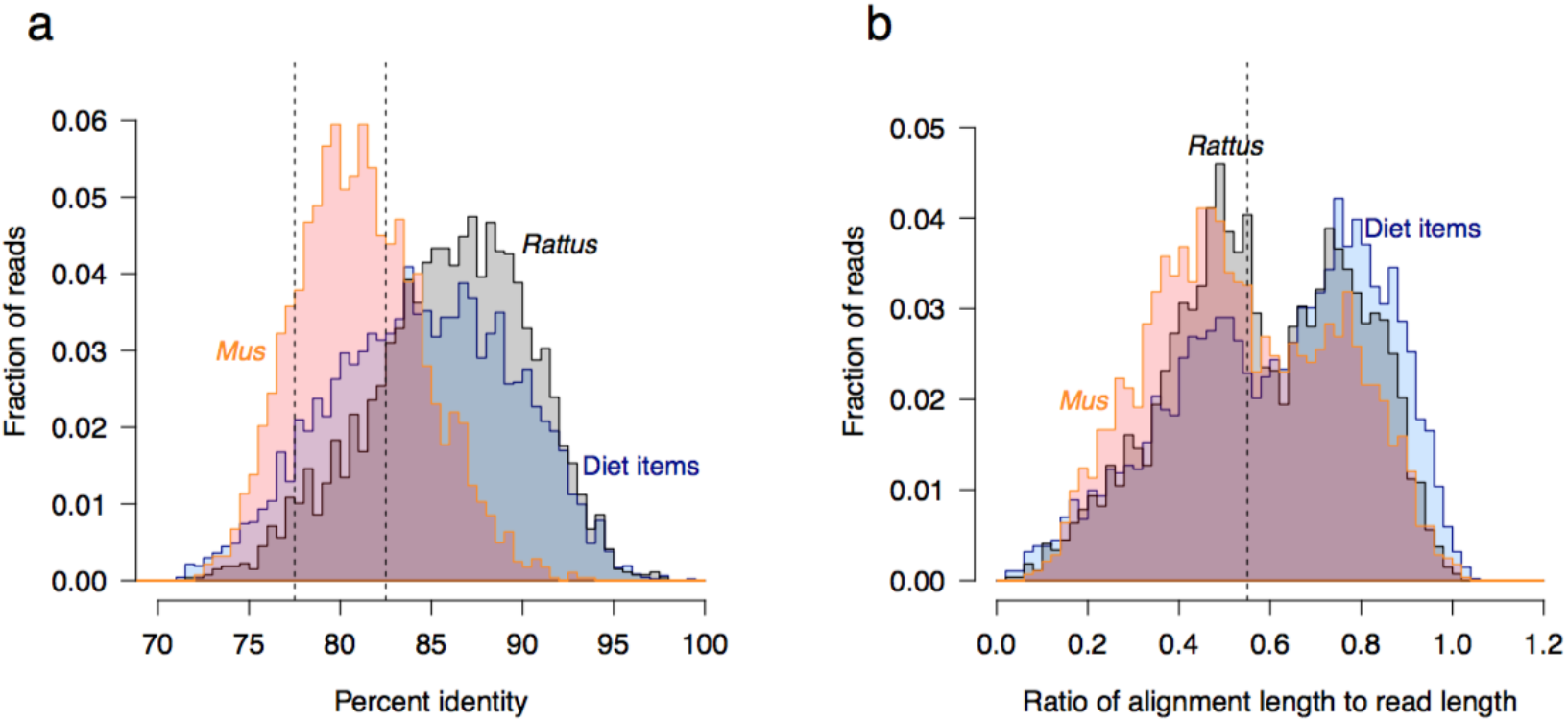
Distributions of percent identity and length for alignments of reads matching *Rattus* (rat), *Mus* (mouse), and diet items. (a) The percent identity for alignments of rat (*Rattus*) and diet items is much higher than for mouse (*Mus*). Histograms of the percent identity of the alignment of the top BLAST hit with the read. *Mus* matches have substantially lower percent identity compared to both *Rattus* and diet items. The dotted lines indicate the cut-offs that we implemented for inferring reads as belonging to a specific genus (above 82.5% identity) or family (above 77.5% identity). **(b) Ratios of alignment lengths to read lengths of rat (*Rattus*) and diet items are higher than for mouse (*Mus*).** This plot is analogous to that in (a). The dotted line indicates the cut-off that we implemented for inferring reads as belonging to a specific genus (above 0.55).

To further investigate alignment characteristics differentiating true positive (*Rattus*) and false positive (*Mus*) assignments, we considered the ratio of alignment length to read length. If the full read aligns, this ratio is one. More generally, higher quality alignments should have higher ratios. Indeed, we found this ratio differed substantially between the *Rattus*- and *Mus*-assigned reads: the median ratio of alignment length to read length was 0.57 for *Rattus* and 0.52 for *Mus* (Fig. 3B).

We used this analysis to select cut-off values for assigning reads as having true positive or false positive taxon assignments. For genus-level assignments, we required at least 82.5% alignment identity, and an alignment length to read length ratio of at least 0.55. These cut-offs exclude 88% of the reads falsely assigned to *Mus*, instead correctly assigning them one taxon level higher, to the Family *Muridae*.

However, this rate of false positive assignments is still relatively high. We thus also quantified the rate of false positive assignment at the level of family. We first identified all reads classified as being from the order *Rodentia.* We then implemented specific cut-offs, requiring reads and database matches to have 77.5% identity, an alignment length to read length ratio of at least 0.1, and a total alignment length of at least 150 bp. With these cut-offs, 97.3% of all reads assigned to the Order *Rodentia* were classified in the likely true positive Family *Muridae* (which contains the genus *Rattus*). The remaining 2.7% were assigned to the family *Cricetidae* (voles and lemmings), except or four reads assigned to *Spalacidae* (mole-rats) (Fig. S2). These are clear false positive assignments, as it is highly unlikely that these families were predated. However, these results establish that by implementing specific cut-off values for alignments, we can ensure a low rate of false positive assignments at the family level, especially for taxa with genomic sequence well-represented in the database.

However, it is possible that for taxa that are not well-represented in databases, read classification will be less accurate. If this occurred, we would expect that reads would frequently be assigned to the wrong taxon, resulting in lower percent identities and low alignment lengths for those reads. We thus checked whether diet items had percent identities and alignment lengths similar to the true positive *Rattus* alignments, or instead whether they were more similar to the false positive *Mus* alignments. We found that the majority of diet items had alignment percent identities that overlapped with the *Rattus* reads. Furthermore, the alignment length to read length ratios often exceed those for the *Rattus* reads. This suggests that the diet taxa assignments are often correct down to the level of genus (as the *Rattus*-assigned reads are correct to the level of genus) and are not false positive assignments. However, here we conservatively report diet items at the level of family.

For reads that did not pass the above cut-offs, we placed taxon assignments at the level of order, or used the taxon level assigned by MEGAN. Using these cut-offs, 16% of all reads were classified at the genus level; 71% were classified at the family-level or below; 89% were classified at the order-level or below; and 98% were classified at the phylum-level or below.

Beyond the small fraction of false positive family assignments within *Rodentia* (discussed above), a small fraction of clear false positives remained in other orders. Again, most had short alignments (Table S4). The exception to this were three reads from two rats matching *Buthidae* (scorpions), which had alignment lengths of 762 bp, 664 bp, and 298 bp with identities of 83%, 88%, and 79%, respectively. It is unlikely these are true positives, and instead we hypothesise that these rats predated harvestmen (*Opiliones*), a closely related sister taxon within *Arachnida*, but lacking significant amounts of genomic data. Despite the presence of these false positive taxa, we did not further increase the stringency of our filters, suggesting that we have a small rate of false positive inference at the family level (2.7% in the case of *Muridae*), and most likely similar rates for other family assignments.

### Identification of diet

Within each rat, we identified a wide variety of plant, animal, and fungal orders, ranging from two to 25 orders per rat (mean 8.7; Fig. 4). In total, we identified taxa from 68 different Families, 55 different Orders, 15 different Classes, and eight different Phyla (Fig. 5). In this analysis, we consider all non-host, non-bacterial reads as potential diet items, although not all are necessarily so. Plants were the primary diet item, with the largest fraction of rats consuming four predominant orders: *Poales* (grasses), *Fabales* (legumes), *Arecales* (palms), and *Araucariales* (podocarps). The dominance of plant matter (fruits and seeds) in rat diets has been established previously (57,58). Animal taxa made up a smaller component of each rat’s diet, with *Insecta* dominating: *Hymenoptera* (bees, wasps, and ants), *Coleoptera* (beetles), *Lepidoptera* (moths and butterflies), *Blattodea* (cockroaches), *Diptera* (flies), and *Phasmatodea* (stick insects). In addition, *Stylommatophora* (slugs and snails) were present in substantial numbers (Fig. 5A and 5B). Fungi were only a small component of the rats’ diet, although several orders were present: *Sclerotiniales* (plant pathogens), *Saccharomycetales* (budding yeasts), *Mucorales* (pin molds), *Russulales* (brittlegills and milk-caps), and *Chytotheriales* (black yeasts). Finally, for many rats, a substantial proportion of the stomach contents were parasitic worms (primarily *Spirurida* (nematodes) and *Hymenolepididae* (tapeworms)).

**Fig. 4.**
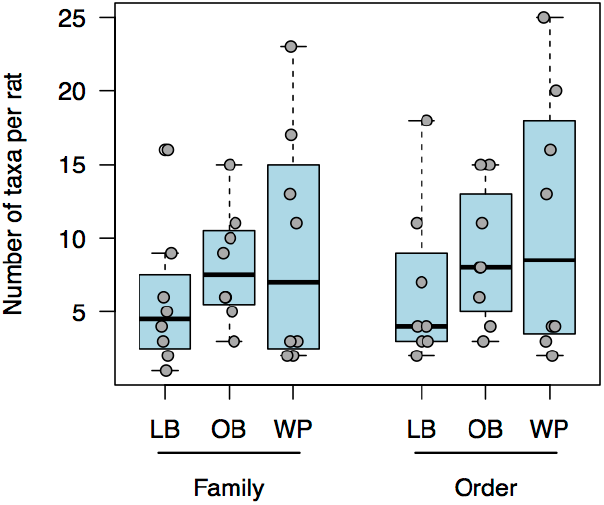
Numbers of taxa in individual rats. Each boxplot indicates the range of families (left boxes) or orders (right boxes) consumed by each rat in each location (OB: Okura Bush, native bush; LB: Long Bay Park, restored wetland; WP: Waitakere Park, native forest). The numbers for individual rats (eight per location) are plotted in grey.

**Fig. 5.**
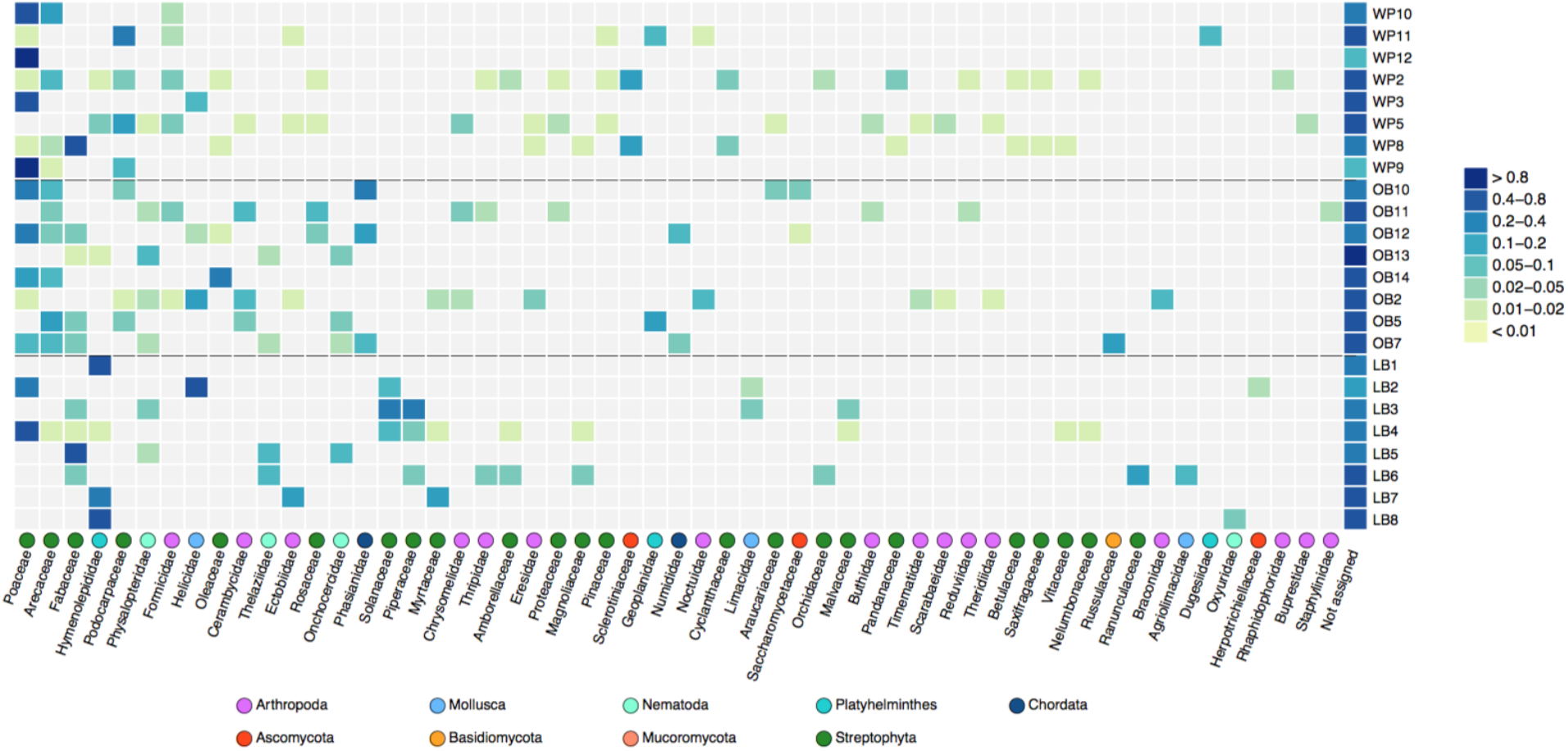
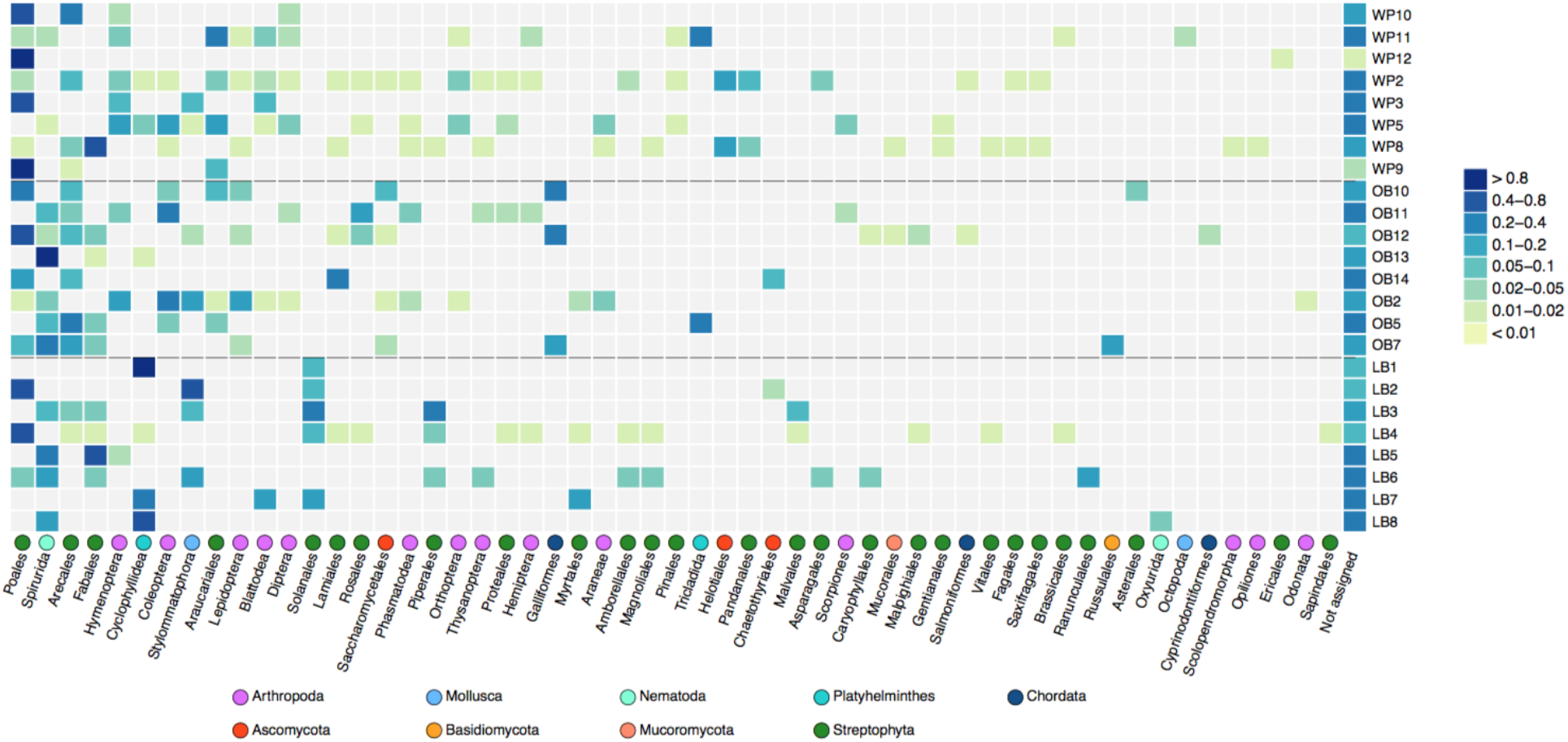
Proportions of taxa in the diets of individual rats. (a) Reads assigned to taxa at the family and (b) order level. The rows correspond to a single rat, with the proportions of reads for that rat assigned to each family or order indicated in shades of blue and yellow. Reads that were not assigned to a specific family or order are indicated at the right side of the figure. The families and orders have been sorted so that the most common diet components appear on the left. Only the 55 most common families are shown. Note that the color gradations presented on the scale are not linear.

Due to our metagenomic approach, the fraction of each element of the rats’ diets is distorted by biases in genomic databases: whole genome data exists for only a few taxa, while mtDNA, rDNA, metabarcode loci, and microsatellite sequence data are present in the database for many animal and plant genera. To quantify database-driven bias, for each taxon we determined the fraction of hits that mapped to mtDNA, rDNA, microsatellites, or EST libraries (we refer to this as *non-genomic*), versus the fraction of hits that mapped to DNA sequences annotated as arising from genome sequencing projects). We expect that for animals with sequenced genomes, these two fractions should be approximately constant, largely determined by the relative amounts of mtDNA and nuclear DNA in the cell. For animals without sequenced genomes, there should be considerably more hits to mtDNA, plastid, rDNA, and microsatellites (non-genomic sequence), and few (if any) genomic hits. By comparing these fractions for taxa with complete genome sequences in the database to taxa without complete genomes we aimed to assess the effects of this bias.

We first considered only animals with sequenced genomes in the database. For these, we found that the fraction of reads that mapped to non-genomic sequence (mtDNA, rDNA, microsatellite, plastid, and EST library) ranged from 0% (*Coturnix* and *Numida*) to 27% (*Rattus*) to 39% (*Gallus*) (Fig. 6). This is a considerable range, and we hypothesise that variation in the fraction of non-genomic reads is due to the type of tissue sequenced. For *Rattus* the sequenced tissue was primarily stomach muscle, which has a relatively high fraction of mtDNA, explaining the large fraction of reads mapping to mtDNA.

**Figure 6.**
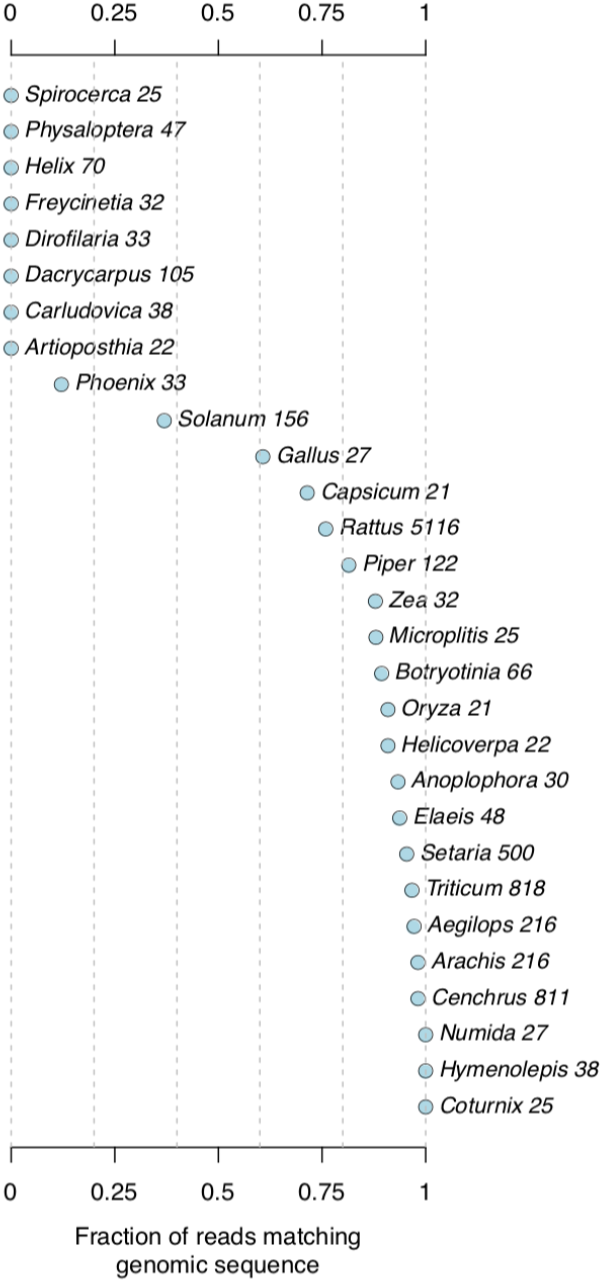
Fractions of reads matching genomic and non-genomic sequence for the best BLAST hit of each read. For the species with complete genomes, the fraction of reads matching genomic sequence ranges from 40% (*Solanum*) to 100%. This large range is likely due to the tissue from which the DNA was isolated. For example, muscle tissue has a higher fraction of mtDNA to nuclear DNA than egg. For species without fully sequenced genomes, this fraction ranges from 0% to 20% (*Phoenix*, which has a small amount of genomic data present in the database).

For plants with sequenced genomes, the fraction of reads matching non-genomic sequence (mostly mtDNA, plastid, and rDNA) was generally lower: between 2% (*Cenchrus*) and 12% (*Zea*). Thus, on average, for animals with sequenced genomes present in the database, approximately 30% of all reads mapped to non-genomic sequences; for plants, approximately 5% mapped to non-genomic sequences.

In contrast, for taxa with little or no genomic sequence in the database, the vast majority of matches were non-genomic (mtDNA, plastid, rDNA, or microsatellite loci): 90% of *Phoenix* (date palm) hits; all *Helix* (snail); and all *Rhaphidophorae* (cave weta) hits. All *Arthurdendyus* (the endemic New Zealand flatworm) hits were solely to rDNA loci.

This overrepresentation of non-genomic sequence is in strong contrast to animals that do have sequenced genomes. In that case we expect only 30% of all reads should map to non-genomic sequence. Thus, these results indicate that for animal taxa with little or no genomic sequence data, we have underestimated the actual number of sequences from that taxon by approximately two-to three-fold. For plant taxa with little or no genomic sequence data, we have likely underestimated read abundance by approximately 20-fold.

After classification of the reads at the genus, family, and order level, further examination of the data suggested that specific taxa were overrepresented in the diets of rats from particular locations. For example, six out of eight rats from the native estuarine bush habitat (OB) consumed *Arecaceae*, while only one in the restored wetland area (LB) did. All three rats that consumed *Phaseanidae* were from the native estuarine habitat (OB). All five rats that consumed *Solanales* were from the restored wetland area. These patterns suggested that it might be possible to use diet components alone to pinpoint the habitat from which each rat was sampled.

### nMDS and CAP analysis by location

In order to determine if diet composition of the rats differed consistently between locations, we first performed an unconstrained analysis using non-metric multidimensional scaling (nMDS) on taxa assigned at the family level. The input for the nMDS was the dissimilarity matrix (Bray-Curtis distance of diet at the family level). NMDS uses rank-based distances to cluster samples that are most similar.

The family-level unconstrained ordination (nMDS) showed no obvious grouping of rats with respect to the locations (Fig. 7A), indicating that locations did not correspond to the predominant axes of variation among the diets. We next performed a constrained ordination method, Canonical Analysis of Principal coordinates (CAP, see Methods). CAP identified axes of variation, if any, that distinguish the diets of rats from different locations (Fig. 7B). We found that the CAP axes correctly classified the locations of 19 out of 24 (79%) rats using a leave-one-out procedure. The families having the largest correlations with the first two principal coordinates, and thus most responsible for the separation between groups, were primarily plants: *Arecaceae*, *Podocarpaceae*, *Piperaceae*, and *Pinaceae*. In addition, insect groups (*Cerambycids* and *Formicids*) and birds (*Phaseanidae* and *Numididae*) played a role (Fig. 7C).

**Fig. 7.**
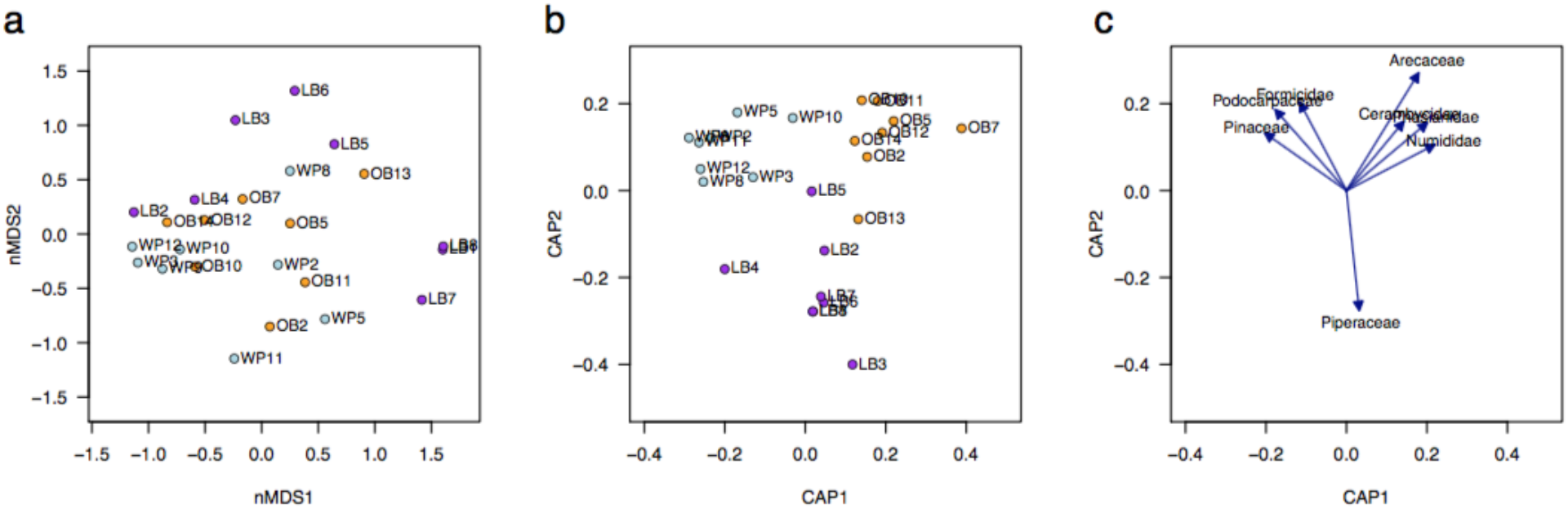
Unconstrained nMDS (a) and constrained CAP (b) ordinations of the diets of rats from three locations. Both ordinations were based on Bray-Curtis dissimilarities of square root transformed proportions of reads attributed to each family. The locations were a native estuarine bush (OB, orange); a restored marine wetland (LB, purple); and a native forest (WP, light blue). The CAP ordination is repeated in panel **(c)** as a biplot with the rats omitted to show the Pearson correlations between families and the first two CAP axes. The eight families with the strongest correlations are shown, indicating the taxa associated with each location.

The families driving similarity within the three locations (i.e., those that had the greatest within-location SIMPER scores, see Methods) varied among locations. LB had average Bray-Curtis within-location similarity of 13%; mostly attributable to *Hymenolepidae* (accounting for 51% of the within-group similarity), *Solanaceae* (11%), and *Fabaceae* (11%). The average similarity for OB was 21%, with the greatest contributing taxa being *Arecaceae* (33%), *Poaceae* (23%), *Fabaceae* (9%), and *Phasianidae* (8%). The average similarity for WP was 24%, with the greatest contributing taxa being *Poaceae* (72%) (Table S4).

## Discussion

### Accuracy and sensitivity

Here we have shown that using a simple metagenomic approach with error-prone long reads allows rapid and accurate classification of rat diet components (approximately 2.7% error in taxon assignments at the family-level). We expect that this technique can be used to infer diet for a wide variety of animal and sample types, including samples that use less invasive collection methods, such as fecal matter. The sensitivity of this approach will likely improve as the accuracy and yield of ONT sequencing continues to increase. The analysis here is based on less than 200,000 reads from two flow cells. Current yields for similar read length distributions are in excess of ten million reads per flow cell. As ONT sequencing accuracy improves (currently just above 96% modal accuracy), this will also increase the applicability of this method. This is illustrated by the fact that the fraction of reads yielding BLAST hits increased substantially for higher quality reads, approaching 40% for high quality reads in our dataset (reads with greater than 92.5% accuracy, Fig. 2B).

As the species sampling of genomic databases increases (59), the taxon-level precision of this method will improve. Given the current rate of genomic sequencing, with careful sampling, the vast majority of multicellular plant and animal families (and even genera) will likely have at least one type species with a sequenced genome within the next decade. Continued advancement in sequence database search algorithms as compared to current methods (23,24,60) should considerably decrease the computational workload necessary to find matching sequences.

### Methodological advantages

As genomic databases become more complete, metagenomic approaches will offer significant advantages due to decreased bias. We found that rats consumed many soft-bodied species (e.g. mushrooms, flat worms, slugs, and lepidopterans) that would be difficult to identify using visual inspection of stomach contents. Achieving data on such a wide variety of taxa (across multiple phyla) would be difficult to quantify using metabarcoding, as there are no universal 18S or COI universal primers capable of amplifying sequences in all these taxa. While it might be possible to use several different primer sets targeted at different phyla or orders, quantitatively comparing diet components across these using sequences amplified with different primer sets is extremely difficult due to differences in primer binding and PCR efficiency. A second advantage of metagenomic sequencing (as has been pointed out previously), is that it allows for “holistic” views of ecology, as it simultaneously profiles parasites, microbiome, and prey species (30).

The ONT-based sequencing method has several unique advantages, and perhaps the most obvious is the accessibility of the platform. Compared to other high throughput sequencing technologies (e.g. Illumina, IonTorrent, or PacBio), there is no initial capital investment required to use the platform. On a per-sample basis, data generation is inexpensive (assuming 12 multiplexed samples, approximately $150 USD per sample, and half this price if reagents are purchased in bulk). Library preparation and sequencing can be extremely rapid, going from DNA sample to sequence in less than two hours (61). Furthermore, the sequencing platform itself is highly portable. Given (1) that ONT-based methods are now similar in cost-per-read (approximately $650 for 10 million reads) than the most accessible Illumina method (MiSeq, approximately $1300 for 20 million reads); and (2) that even marginal increases in read length are likely to significantly improve species identification, we expect that ONT-based methods should soon become useful for routine ecological monitoring of species (62).

Some modifications to our approach might further increase the precision of our ability to infer community composition. Any error-prone long read dataset (i.e. PacBio or ONT) has both short (e.g. 500 bp) and long (e.g. 5000 bp) reads, as well as high quality (e.g. mean accuracy greater than 90%) and low quality (e.g. mean accuracy less than 80%) reads. When inferring community composition, a null expectation is that taxa should be equally represented by long, high quality reads as they are by short, low quality reads. If some taxa are represented only by short, low quality reads, this suggests that these taxa may be false positive inferences. Similarly, the difficulty in correctly mapping short inaccurate reads could be mitigated by weighting the probability of taxon mapping by the number of long, accurate reads that map to certain taxa. Thus, the fact that not all reads are extremely long and accurate does not mean that they cannot all be used to infer taxon presence in metagenomic analyses.

For many diet studies, the aim is to ideally resolve the biomass, or nutritional content, or prey numbers. However, as a caveat to the methods utilised here, and to all sequencing-based methods, the accuracy of estimating these numbers is constrained by the fact that different tissues and different taxa have different amounts of DNA (both nuclear and mitochondrial) per gram of biomass. It is nearly to impossible to fully account for this variation. Regardless, there is considerable utility in using sequencing based approaches for diet assessment, not least because it is one of the few methods that allows the full breadth of the diet to be observed, as illustrated here by the number of different orders we find that rats predate.

### Conclusions

Here we have shown that a rapid error-prone long read metagenomic approach is able to accurately characterise diet taxa at the family level and distinguish between the diets of rats according to the locations from which they were sourced. This information may be used to guide conservation efforts toward specific areas and habitats in which native species are most at risk from this highly destructive introduced predator.

## Methods

### Study Areas

We trapped rats from three locations near Auckland, New Zealand. Each location comprised a different type of habitat: undisturbed inland native forest (Waitakere Regional Parklands, WP); native bush surrounding an estuary (Okura Bush Walkway, OB); and restored coastal wetland (Long Bay Regional Park, LB) (Fig. 8). Snap traps in OB and LB were baited with peanut butter, apple, and cinnamon wax pellets; or bacon fat and flax pellets.

**Fig. 8.**
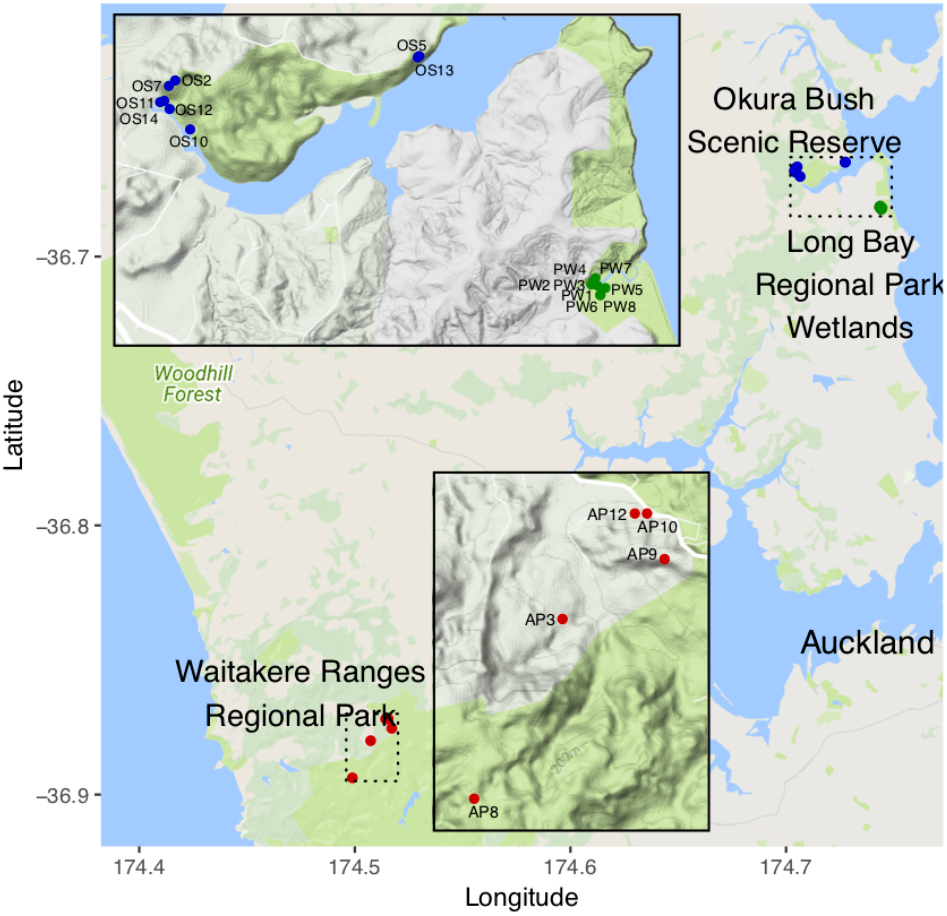
Location of rat sampling sites in the greater Auckland area in the North Island of New Zealand. Each point indicates a trap where one rat was captured, with the colour of the points indicating the three broad locations: the native estuarine bush habitat of Okura Bush (OB), the restored wetland of Long Bay (LB), and the native forest of Waitakere Park (WP). The two insets show the three locations in higher resolution with topographical details. Green indicates park areas. Precise geographical coordinates were only available for five out of eight rats in WP.

Traps in WP were baited with chicken eggs, rabbit meat, or cinnamon scented poison pellets. From 16 November to 16 December 2016, traps were surveyed by established conservation groups at each site every 48 hours. A total of 36 rats were collected from these locations. Three of the rats from WP had poison in their stomachs. However, all three of these were killed by the snap traps, and as such are unlikely to have swallowed any bait. In addition, none were identified as having chicken or rabbit in their diet. The majority of rats collected (34/36) were determined to be male *Rattus rattus* by visual inspection. These 34 rats were selected for further analysis.

### DNA Isolation

Within 48 hours of trapping, rats were stored at either −20°C or −80°C until dissection. We dissected out intact stomachs from each animal and removed the contents. After snap freezing in liquid nitrogen, we homogenised the stomach contents using a sterile mini blender to ensure sampling was representative of the entire stomach.

We purified DNA from 20 mg of homogenised stomach contents using the Promega Wizard Genomic DNA Purification Kit, with the following modifications to the Animal Tissue protocol: after protein precipitation, we transferred the supernatant to a new tube and centrifuged a second time to minimise protein carryover. The DNA pellet was washed twice with ethanol. These modifications were performed to improved DNA purity. We rehydrated precipitated DNA by incubating overnight in molecular biology grade water at 4°C and stored the DNA at −20°C. DNA quantity, purity, and quality was ascertained by nanodrop and agarose gel electrophoresis. The DNA samples were ranked according quantity and purity (based on A260/A280 and secondarily, A230/A280 ratios). The eight highest quality DNA samples from each of the three locations were selected for sequencing. We did no size selection on the purified DNA.

### DNA Sequencing

Sequencing was performed on two different dates (24 January 2017 and 17 March 2017) using a MinION Mk1B device and R9.4 chemistry. For each sequencing run, DNA from each rat was barcoded using the 1D Native Barcoding Kit (Barcode expansion kit EXP-NBD103 with sequencing kit SQK-LSK108) following the manufacturer’s instructions. This included an AMPure bead purification step to remove adaptors, which also likely removed very short reads (less than 200 bp; see Fig. 1A). Twelve samples were pooled and run on each flow cell, for a total of 24 individual rats. The flow cells had 1373 active pores (January 2017) and 1439 active pores (March 2017). Sequencing was performed using local base calling in MinKnow v1.3.25 (January) or MinKnow v1.5.5 (March), but for the analysis present here both runs were re-basecalled after data collection using Albacore 2.2.7 with demultiplexing performed in Albacore and filtering disable*d (optio*ns *--barcoding --disable_filtering*).

### Sequence classification

All sequences were BLASTed (blastn v2.6.0+) against a locally compiled database consisting of the combined NCBI other_genomic and nt databases (downloaded on 13^th^ June 2018 from NCBI). Default blastn parameters were used (match 2, mismatch −3, gapopen −5, gapextend −2), and only hits with an e-value of 1e-2 or less were saved. Due to the predominance of short indels present in nanopore sequence data, we used an initial set of basecalled data to test whether changing these default penalties affected the results (gapopen −1, gapextend −1). We found that these adjusted parameters did not qualitatively change our results.

We assigned sequence reads to specific taxon levels using MEGAN6 (v.6.11.7 June 2018) (41). We only used reads with BLAST hits having an e-value of 1×10^−20^ or lower (corresponding to a bit score of 115 or higher given the databases we used) and an alignment length of 100 base pairs or more. To assign reads to taxon levels, we considered all hits having bit scores within 20% of the bit score of the best hit (MEGAN parameter Top Percent).

### Multivariate analyses

Multivariate analyses were done using the software PRIMER v7 (42). The data used in the multivariate analyses were in the form of a sample-(i.e. individual rat) by-family matrix of read counts. All bacteria, rodent, and primate families were removed. The majority of rodent hits were to rat and mouse, resulting from the rats’ own DNA (see below). The majority of the primate hits (32 in total) were assigned to Hominidae (19), which likely resulted from sample contamination (Table S3).

The read counts were converted to proportions per individual rat by dividing by the total count for each rat, to account for the fact that the number of reads varied substantially among rats (43). The proportions were then square-root transformed so that subsequent analyses were informed by the full range of taxa, rather than just the most abundant families (44). We then calculated a matrix of Bray-Curtis dissimilarities, which quantified the difference in the gut DNA of each pair of rats based on the square-root transformed proportions of read counts across families (43).

We used unconstrained ordination, non-metric multidimensional scaling (nMDS) applied to the dissimilarity matrix to examine the overall patterns in the diet composition among rats. To assess the degree to which the diet compositions of rats were distinguishable among the three locations, we applied canonical analysis of principal coordinates (CAP) (45) to the dissimilarity matrix. CAP is a constrained ordination which aims to find axes through multivariate data that best separates *a priori* groups of samples (in this case, the groups are the locations from which the rats were sampled); CAP is akin to linear discriminant analysis but it can be used with any resemblance matrix. The out-of-sample classification success was evaluated using a leave-one-out cross-validation procedure (45).

We used Similarity Percentage (SIMPER; (46)) to characterise and distinguish between the locations. This allowed us to identify the families with the greatest percentage contributions to (1) the Bray-Curtis similarities of diets within each location (Table S5) and (2) the Bray-Curtis dissimilarities between each pair of locations (Table S6).

## Supporting information

Datafile S1, table of taxon assignments

Datafile S2, filtered table of taxon assignments

## Declarations

### Ethics approval

Sample collection was performed under (Auckland Council Permit to Undertake Research WS1064).

### Availability of data and materials

Sequence data are available in the SRA archive (accession number PRJEB27647).

### Authors’ contributions

WP, JD, NF, and OS conceived the project. WP performed the stomach dissections. WP and NF optimised the genomic DNA isolation and library preparation. NF performed the nanopore sequencing. GB and OS processed and performed quality control on the sequencing data. WP and OS performed the sequence classification. WP, AS, NF, and OS analysed the data. WP, NF, AS, and OS wrote the paper, with input from all authors.

## Acknowledgements

Thanks to Friends of Okura Bush, Mary Stewart from Auckland Council, and Gillian Wadams and the volunteers at the Waitakere Ranges for collecting rat samples and aiding in rat species identification.

---

**Table S1.** Read numbers and total base pairs for each barcode in the January 2017 sequencing run.

| Sample | Total reads | Total Mbp | Mean length |
| --- | --- | --- | --- |
| OB2 | 19907 | 14.62 | 734 |
| WP11 | 10164 | 9.63 | 947 |
| WP5 | 8237 | 6.78 | 823 |
| LB7 | 7548 | 7.04 | 933 |
| OB13 | 3644 | 3.63 | 995 |
| WP9 | 2954 | 2.4 | 814 |
| OB5 | 2850 | 2.06 | 721 |
| WP8 | 2801 | 2.32 | 827 |
| LB6 | 2531 | 1.6 | 632 |
| OB7 | 2473 | 1.87 | 756 |
| LB5 | 1641 | 1.16 | 705 |
| LB3 | 1554 | 0.99 | 636 |
| None | 16673 | 13.01 | 781 |
| Total | 82977 | 67.1 | 809 |

**Table S2.** Read numbers and total base pairs for each barcode in the March 2017 sequencing run.

| Sample | Total reads | Total Mbp | Mean length |
| --- | --- | --- | --- |
| LB1 | 17820 | 9.21 | 517 |
| LB8 | 16923 | 13.13 | 776 |
| WP2 | 10511 | 7.00 | 666 |
| LB4 | 8684 | 4.92 | 567 |
| OB11 | 5689 | 3.40 | 598 |
| WP10 | 1563 | 0.99 | 633 |
| OB12 | 1479 | 0.89 | 604 |
| WP12 | 1309 | 0.78 | 596 |
| LB2 | 1127 | 0.76 | 676 |
| WP3 | 637 | 0.73 | 1141 |
| OB14 | 541 | 0.37 | 683 |
| OB10 | 435 | 0.24 | 555 |
| None | 29432 | 21.33 | 725 |
| Total | 96150 | 63.75 | 663 |

**Table S3.** Characteristics of alignments for reads assigned to the Primate family. Many are both long and have high identity, suggesting that they are not false positive assignments, but contamination.

| Rat | Read length | Mean read accuracy | % ID | Alignment length | Genus |
| --- | --- | --- | --- | --- | --- |
| WP10 | 428 | 0.91 | 94.0 | 314 | Homo |
| WP10 | 782 | 0.92 | 93.8 | 657 | Homo |
| OB5 | 365 | 0.90 | 93.2 | 249 | Homo |
| LB3 | 515 | 0.95 | 92.6 | 462 | Homo |
| WP10 | 510 | 0.90 | 92.0 | 460 | Homo |
| WP10 | 704 | 0.90 | 91.1 | 587 | Homo |
| WP10 | 467 | 0.89 | 90.3 | 402 | Homo |
| WP10 | 494 | 0.89 | 89.7 | 388 | Homo |
| WP10 | 339 | 0.88 | 89.6 | 269 | Homo |
| WP10 | 446 | 0.88 | 89.6 | 326 | Homo |
| WP10 | 327 | 0.91 | 89.1 | 210 | Homo |
| OB5 | 415 | 0.89 | 88.8 | 277 | Homo |
| OB5 | 561 | 0.86 | 88.7 | 257 | Homo |
| WP11 | 434 | 0.86 | 88.7 | 301 | Homo |
| WP10 | 486 | 0.88 | 88.2 | 365 | Homo |
| WP10 | 613 | 0.90 | 88.2 | 526 | Homo |
| WP10 | 563 | 0.87 | 88.2 | 457 | Homo |
| WP10 | 1373 | 0.91 | 87.7 | 1337 | Homo |
| WP11 | 526 | 0.91 | 87.5 | 473 | Homo |
| OB14 | 478 | 0.89 | 86.9 | 373 | Homo |
| OB5 | 715 | 0.89 | 86.8 | 673 | Homo |
| WP10 | 475 | 0.87 | 86.7 | 362 | Homo |
| WP10 | 398 | 0.86 | 86.5 | 259 | Homo |
| WP10 | 377 | 0.88 | 85.3 | 251 | Homo |
| WP9 | 558 | 0.87 | 85.2 | 508 | Homo |
| WP10 | 429 | 0.86 | 84.8 | 276 | Homo |
| WP10 | 322 | 0.81 | 84.5 | 174 | Homo |
| WP8 | 723 | 0.84 | 83.1 | 438 | Homo |
| LB8 | 965 | 0.86 | 80.4 | 245 | Rhinopith<br>ecus |
| LB5 | 3042 | 0.94 | 79.4 | 1018 | Cebus |
| WP2 | 464 | 0.93 | 77.3 | 216 | Homo |
| LB8 | 671 | 0.90 | 73.9 | 406 | Aotus |

**Table S4.**
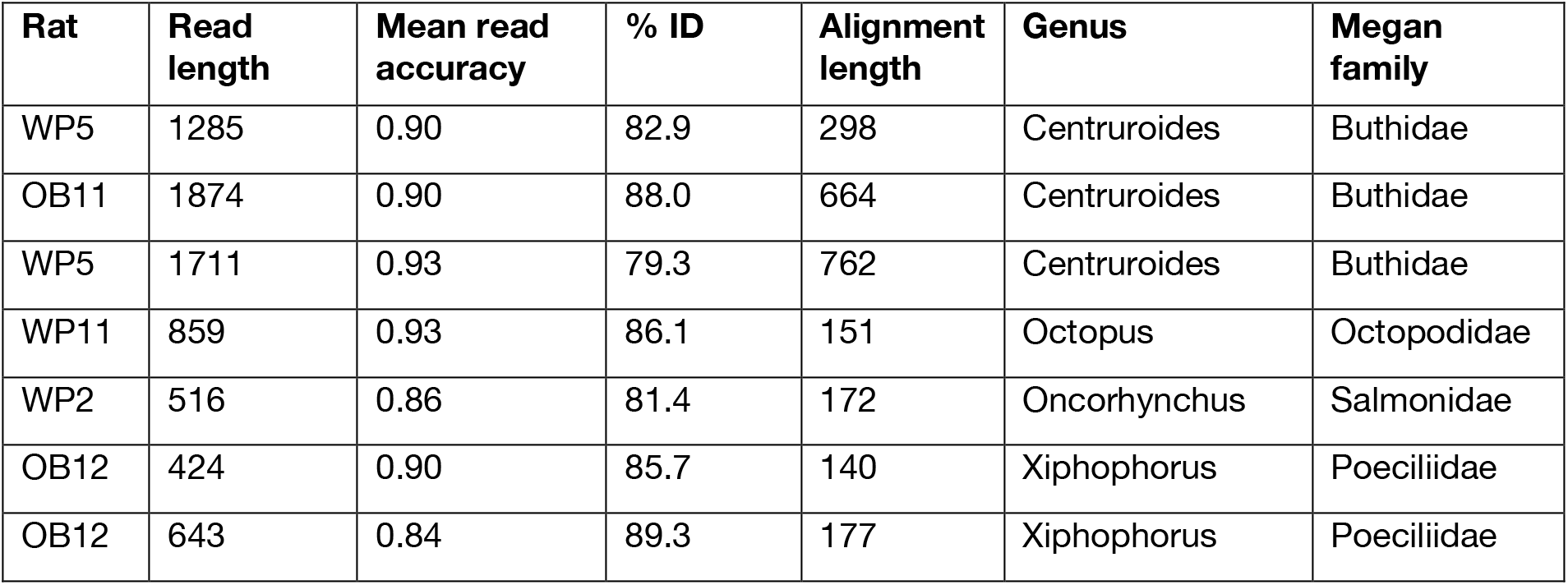
Characteristics of alignments for reads that are likely false positive assignments. Most are short long or have low identity, suggesting that they are false positive assignments. The exception are the reads matching Buthidae, which we hypothesize are due to the rats predation of the sister taxa, harvestmen.

| Rat | Read length | Mean read accuracy | % ID | Alignment length | Genus | Megan family |
| --- | --- | --- | --- | --- | --- | --- |
| WP5 | 1285 | 0.90 | 82.9 | 298 | Centruroides | Buthidae |
| OB11 | 1874 | 0.90 | 88.0 | 664 | Centruroides | Buthidae |
| WP5 | 1711 | 0.93 | 79.3 | 762 | Centruroides | Buthidae |
| WP11 | 859 | 0.93 | 86.1 | 151 | Octopus | Octopodidae |
| WP2 | 516 | 0.86 | 81.4 | 172 | Oncorhynchus | Salmonidae |
| OB12 | 424 | 0.90 | 85.7 | 140 | Xiphophorus | Poeciliidae |
| OB12 | 643 | 0.84 | 89.3 | 177 | Xiphophorus | Poeciliidae |

**Table S5.** SIMPER analysis of family contributions to group similarities.

| Family | Average | Average | Similarity/SD | Percentage | Group |
| --- | --- | --- | --- | --- | --- |
| Hymenolepididae | 3.37 | 6.87 | 0.34 | 51.2 | LB |
| Solanaceae | 1.57 | 1.48 | 0.34 | 11.1 | LB |
| Fabaceae | 1.74 | 1.41 | 0.44 | 10.5 | LB |
| Arecaceae | 2.86 | 7.11 | 1 | 33.4 | OB |
| Poaceae | 2.87 | 4.82 | 0.55 | 22.7 | OB |
| Fabaceae | 1.17 | 1.98 | 0.51 | 9.3 | OB |
| Phasianidae | 1.79 | 1.67 | 0.34 | 7.9 | OB |
| Poaceae | 5.08 | 17.61 | 0.62 | 72.1 | WP |

**Table S6.** SIMPER analysis of family contributions to group dissimilarities.

| Species | Av Abund<br>Group1 | Avg Abund<br>Group2 | Avg.<br>Diss | Diss/SD | % contrib | Group<br>1 | Group2 |
| --- | --- | --- | --- | --- | --- | --- | --- |
| Poaceae | 1.95 | 5.08 | 15.15 | 1.04 | 16.74 | LB | WP |
| Poaceae | 2.87 | 5.08 | 11.29 | 1.26 | 13.78 | OB | WP |
| Hymenolepididae | 3.37 | 0.48 | 10.8 | 0.73 | 11.93 | LB | WP |
| Hymenolepididae | 3.37 | 0.29 | 9.37 | 0.79 | 10.32 | LB | OB |
| Poaceae | 1.95 | 2.87 | 8.37 | 1.1 | 9.22 | LB | OB |
| Arecaceae | 0.05 | 2.86 | 6.99 | 1.41 | 7.7 | LB | OB |
| Arecaceae | 2.86 | 1.31 | 5.92 | 1.29 | 7.23 | OB | WP |
| Fabaceae | 1.74 | 1.05 | 6.14 | 0.67 | 6.78 | LB | WP |
| Podocarpaceae | 0 | 2.38 | 5.34 | 0.83 | 5.9 | LB | WP |
| Podocarpaceae | 0.71 | 2.38 | 4.82 | 0.99 | 5.88 | OB | WP |
| Fabaceae | 1.74 | 1.17 | 4.87 | 0.81 | 5.37 | LB | OB |
| Fabaceae | 1.17 | 1.05 | 4.31 | 0.84 | 5.26 | OB | WP |

**Datafile S1.** Table of read BLAST hits and assigned MEGAN taxa with with reads reclassified at the family or order level by filtering on read length to alignment length ratio and percent identity.

**Datafile S2.** Table of read BLAST hits and assigned MEGAN taxa with no filters applied.

**Fig S1.**
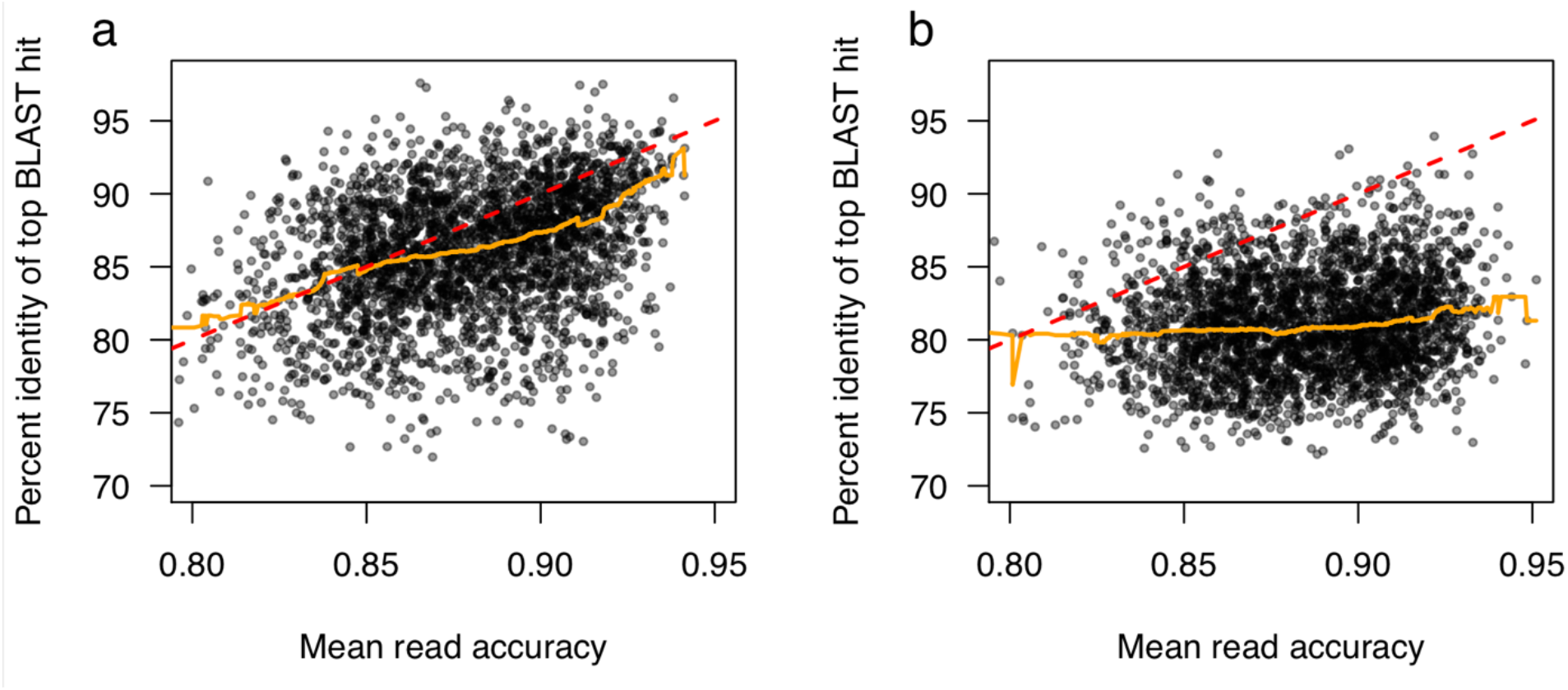
Correlation of read accuracy with alignment characteristics. Only rat reads exhibit a clear positive relationship between accuracy and percent identity. (A) indicates the relationship for reads assigned to *Rattus* and (B) for *Mus*. The orange line indicates a running median; the red dotted line is the y=x line, which is expected if accuracy corresponds exactly to percent identity.

**Fig S2.**
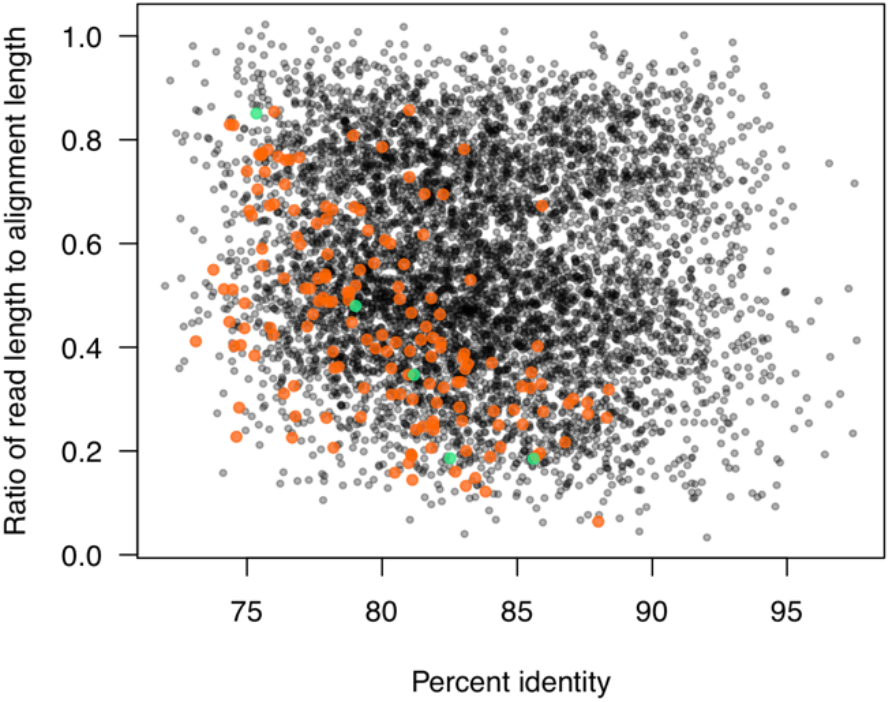
Alignment characteristics of true positive and false positive taxon assignments at the family level. False positive taxon assignments (*Cricetideaa*, orange, and *Spalacidae*, green) have lower percent identity and shorter alignment lengths than true positive taxon assignments (*Muridae*, black). Only a single false positive taxon assignment has a read length to alignment length ratio greater than 0.5 and a percent identity greater than 85%. This suggests that with further filters or methodologies (e.g. decision tree analysis using different read and alignment characteristics) could, if necessary, decrease false positive rates even further.

